# Quantification of glutathione with high throughput live-cell imaging

**DOI:** 10.1101/2023.07.11.548586

**Authors:** Xiaoli Qi, Jianwei Chen, Xiqian Jiang, Dong Lu, Xin Yu, Hanfeng Lin, Erika Y. Monroy, Meng C. Wang, Jin Wang

## Abstract

Reduction oxidation (redox) reactions are central in life and altered redox state is associated with a spectrum of human diseases. Glutathione (GSH) is the most abundant antioxidant in eukaryotic cells and plays critical roles in maintaining redox homeostasis. Thus, measuring intracellular GSH level is an important method to assess the redox state of organism. The currently available GSH probes are based on irreversible chemical reactions with glutathione and can’t monitor the real-time glutathione dynamics. Our group developed the first reversible reaction based fluorescent probe for glutathione, which can measure glutathione levels at high resolution using a confocal microscope and in the bulk scale with a flow cytometry. Most importantly it can quantitatively monitor the real-time GSH dynamics in living cells. Using the 2^nd^ generation of GSH probe, RealThiol (RT), this study measured the GSH level in living Hela cells after treatment with varying concentrations of DL-Buthionine sulfoximine (BSO) which inhibits GSH synthesis, using a high throughput imaging system, Cytation™ 5 cell imaging reader. The results revealed that GSH probe RT at the concentration of 2.0 µM accurately monitored the BSO treatment effect on GSH level in the Hela cells. The present results demonstrated that the GSH probe RT is sensitive and precise in GSH measurement in living cells at a high throughput imaging platform and has the potential to be applied to any cell lines.

## 1. Introduction

### 1.1 Redox reaction homeostasis in health and diseases

Reduction oxidation (redox) reactions are central processes in life maintenance and altered redox state is associated with a spectrum of human ailments, including inflammation and autoimmune diseases, cardiovascular diseases, cancer, and neurodegenerative diseases[1]–[4]. The redox homeostasis is maintained by the balance between oxidants and antioxidants. Reactive oxygen species are continuously generated by aerobic cells and eliminated through antioxidant systems[5]–[7]. Glutathione (GSH) is the most abundant antioxidant in eukaryotic cells and its typical intracellular concentrations range 1 to10 mM[8]. Glutathione is a tripeptide carrying an active thiol group which can be oxidized to form a disulfide bond between two molecules of GSH (GSSG). The reduced (GSH) and the oxidized (GSSG) form of GSH play critical roles in maintaining redox homeostasis in organisms[9]. Besides serving as redox buffer, glutathione also regulates protein functions through S-glutathionylation[10], [11], and acts as a signaling molecule to directly activate gene expression[12]. These important functions are dynamically regulated by the intracellular concentration and distribution of GSH.

### 1.2 Brief history of GSH probe development

Approaches of measuring intracellular GSH have evolved over time and a variety of GSH probes have been developed which include small molecule- and fluorescent protein-based probes[13]. As a small molecule GSH probe, the Ellman’s reagent is still being widely used in GSH measurement[14]. Its poor cell permeability, however, makes it only be used for thiol group detection in pure solutions, blood samples, and extracted tissue samples or cell lysates. Biamine-based probes, such as monobromobimane and monochlorobimane, are non-fluorescent and cell-permeable GSH probes. When incubated with cells, they readily enter cells and form fluorescent complexes which can be detected fluorometrically[15]. Redox sensitive fluorescent proteins (roFPs) measure the glutathione redox potential at a spatial and temporal resolution in living cells based on redox imaging^16^. Since the changes of redox potential in cells could be due to shifts in the [GSH]:[GSSG] ratio or changes in total GSH concentration[17], roFPs could not be considered as GSH-specific probes. The major limitation of the small molecule based GSH probes, the fluorescent protein based probes, and many other reported probes is that these probes are based on irreversible chemical reactions with glutathione, which renders these probes incapable of monitoring the real-time glutathione dynamics. To tackle this problem, we developed the first reversible reaction based fluorescent probe-ThiolQuant Green (TQG)-for glutathione[18]. Moving forward with further optimization of some key properties of the TQG probe, we have already reported the 2^nd^ generation of GSH probe which was designated as RealThiol (RT) and was to be used in this report[17].

### 1.3 Applications of RealThiol in living cells

RT shows ratiometric fluorescence responses with a wide dynamic range when reacting with GSH. RT and its GSH adduct (RT-GSH) shows fluorescence maxima at 562 and 487 nm with excitation wavelengths at 488 and 405 nm, respectively[17]. GSH concentration is calculated as the fluorescence intensity ratios with excitation wavelengths at 405 and 488 nm (F405/F488), which covers the physiological GSH concentration range 1–10 mM. As reported by our group[17], the GSH probe RT has a fast forward and backward reaction kinetics, which enables real-time monitoring of GSH dynamics in living cells. And only micromolar to sub-micromolar RT probe is needed for staining in cell-based experiments, which induces minimal perturbation to GSH level in cells. In addition, the high-quantum-yield coumarin fluorophore moiety minimizes background noise. RT can not only perform high resolution measurements of glutathione levels in single cells using a confocal microscope, but also be applied in flow cytometry to perform bulk measurements. And most importantly this GSH probe can quantitatively monitor the real-time GSH dynamics in living cells[17], [19]. This report will further expand the applications of RT in living cells and explore its capability in GSH measurement at a high throughput imaging platform.

## 2. Material and Methods

### 2.1. Material

1. Glutathione probe RT-AM (Kerafast, Cat. No. EBY001)
2. HeLa cells (ATCC, Cat. No. CCL-2)
3. DMEM (Thermo Fisher Scientific, Cat. No. 10569044)
4. Opti-MEM with reduced FBS and without phenol red (Life Technologies, Cat. No. 11058021).
5. Fetal bovine serum (FBS) (Thermo Fisher Scientific, Cat. No. 10099141)
6. Penicillin streptomycin 10,000 U/mL (Fisher Scientific, Cat. No. 15140122)
7. Trypan blue solution 0.4% (Sigma, Cat. No. 93595, CAS: 72-57-1)
8. Trypsin-EDTA 0.25% (Thermo Fisher Scientific, Cat. No. 25200056)
9. DL-Buthionine sulfoximine (BSO) (Sigma, Cat. No. 19176, CAS: 5072-26-4)
10. Anhydrous DMSO (Sigma, Cat. No. D8418, CAS: 67-68-5)

### 2.2. Equipment and lab wares

1. Cytation™ 5 cell imaging multi-mode reader (BioTek® Instruments, Inc.)
2. Fluorescence filter cube for GSH probe RT, Excitation 469/35 nm, Emission 550/49 nm, Mirror 497 nm
3. Fluorescence filter cube for RT-GSH adduct, Excitation 377/50 nm, Emission 482/25 nm, Mirror 409 nm
4. GSH probe RT LED cube, λ = 465 nm
5. RT-GSH adduct LED cube, λ = 365 nm
6. Cell-culture CO2 incubator, Symphony 5.3 A (VWR)
7. Invitrogen™ Countess™ automated cell counter (Thermo Fisher Scientific Inc.)
8. T-75 flask (Corning, Cat. No. 353810)
9. 96-well tissue culture plate (Corning, Cat. No. 3598)

### 2.3. Software for image capture and data analysis

1. Gene5 version 3.08 (BioTek® Instruments, Inc.)
2. GraphPad Prism 9 (GraphPad Software, Inc.)

### 2.4. Method

Hela cells were maintained in DMEM complete medium and were seeded to 96-well cell culture plates before experiment. After recovering from handling, the cells were treated with serially diluted BSO solutions for 48 h.

Then the cells were stained with GSH probe RT-AM and imaged with a Cytation ™ 5 cell imaging reader. The images were analyzed with the Gene5 software (Fig.1 for flow chart).

**Fig. 1.**
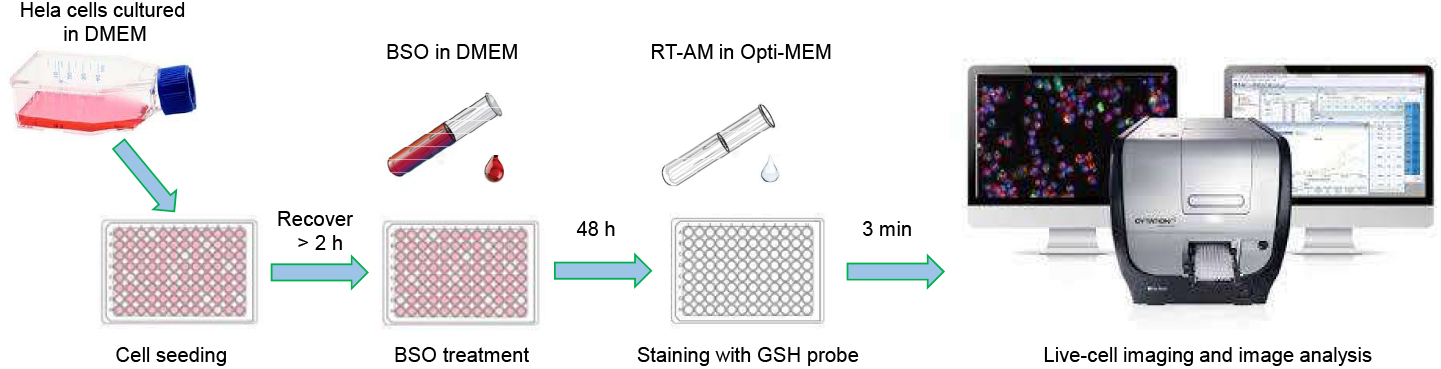
Flow chart for glutathione quantification with high throughput live-cell imaging. Hela cells maintained in DMEM complete medium were seeded to 96-well cell culture plates. After recovering from handling, the cells were treated with serially diluted DL-Buthionine sulfoximine (BSO) solutions for 48 h. Then the cells were stained with RealThiol (RT) probe and were imaged with a Cytation ™ 5 cell imaging reader. The images were analyzed with the software Gene5 version 3.08.

#### 2.4.1 Cell culture and treatment

1. HeLa cells were routinely maintained and cultured in a T-75 flask in DMEM medium supplemented with 10% FBS and 1% penicillin streptomycin antibiotic in a 37 °C incubator with 5% CO2. The typical seeding density was 1 × 10^6^ cells per T-75 flask in 10 mL DMEM complete medium and cells were passaged regularly twice a week. Cells used for tests were typically within 2 months of culture and the passage number is less than 20.
2. For BSO treatment, 2 × 10^3^ HeLa cells were plated in 100 µL of DMEM complete medium in wells of 96-well tissue culture plates, and were allowed to recover for at least 2 hours in an incubator before treatment.
3. BSO stock solution (100 mM) was prepared by dissolving BSO powder in DI water. BSO working solution was prepared by diluting the stock solution with the DMEM complete medium. BSO stock solution can be stored at -20°C for 3 months. Make small aliquots if necessary.
4. After recovery, the cells were treated with serially diluted BSO solutions by adding an equal volume of DMEM complete medium containing 2-fold final concentrations of BSO. The final BSO concentrations were from 500 to 0.9 µ? at 2 folds serial dilution.
5. Cells were incubated for 48 h with varying the concentrations of BSO solution.
6. Prepare RT-AM GSH probe.
  a. Prepare the RT-AM stock solution: dissolve the lyophilized RT-AM probe powder in 10 µL of anhydrous DMSO to make a 5mM stock solution and make aliquots. Stock solution is stable at - 80°C for up to 6 months. **Note:** DMSO is hygroscopic and readily to take up water. Water in DMSO can hydrolyze RT-AM. It is highly recommended, therefore, to dissolve RT-AM in anhydrous DMSO.
  b. Prepare the staining solution: dilute the RT-AM stock solution with DMSO at the ratio of 1:24 to get 200 µM staining solution which can be stored at -20°C for 1 week.
  c. Prepare working solution: the working solution was prepared immediately before use by further diluting the staining solution 100 times in the imaging medium Opti-MEM to get a final concentration of 2.0□μM (with 1% DMSO). Apply 50 -100 µL of working solution to each well of cells pretreated with BSO for 48 h. **Note:** RT-AM working solution is easy to hydrolyze at room temperature and needs to be added to cells immediately after preparation.
7. Aspirate the original DMEM culture medium from cells as much as possible and incubate the cells with 50 -100 µL of RT-AM working solution at room temperature for 3 min. Then the cells were imaged with a Cytation™ 5 Imager.

**Note:** Aspirate the original culture medium as much as possible since the phenol red in the medium may increase background during imaging. PBS, however, is generally not suggested for cell wash as the GSH level was observed change after the cells were exposed to PBS for more than 5 min.

#### 2.4.2 Live-cell imaging and image analysis

##### Imager setup

1. Turn on the Cytation™ 5 imaging reader (Imager), the CO2 gas controller (5%) and CO2 line, and the computer installed with the software Gen5 which controls the Imager for image acquiring and processing as well as data reduction of images captured.
2. Set the imaging chamber temperature to 37 °C in Instrument Control.
3. In the Gen5 software, check the system configuration for objectives, LED cubes, and imaging filter cubes. For the goal of high throughput imaging, a low-power objective lens (4 x) was used to image more cells in one imaging field. Select filter cubes for GSH probe RT (Ex/Em 469/550 nm) and RT-GSH adduct (Ex/Em 377/482 nm) and select corresponding LED cubes (λ= 465 nm and λ= 365, respectively). If the selected cubes are not installed, install new filter cubes and LED cubes in the chamber, restart the Imager, and calibrate the imaging system via Auto Calibration” which typically takes 15 to 30 min and should be allowed for enough time ahead of imaging.
4. Plate definition. For the plate used for the first time in the Imager, it is critical to define the plate bottom elevation and thickness for optimal imaging. Setup of plate parameters will enable the Imager to easily find a reference plane and rapidly focus in a high throughput imaging settings. Also, it will protect the objectives and samples.
5. Let the system equilibrate for at least 30 min before image capture. **Note:** Plate definition step may take longer time than expected. Long exposure of cells under lasers will compromise cell health and affect experiment results. It is highly recommended to prepare an extra plate of cells for parameter optimization and imaging setups.

##### Image acquisition

1. Place the cell culture plate in the imaging chamber.
2. Create a new experiment in Task Manager and set up a Standard Protocol.
3. Select the plate type that has been defined in the Image Setup step and select the wells to be imaged.
4. Set up Read steps for the imaging.
5. Select the 4 x objective.
6. Add channels for GSH probe RT and RT-GSH adduct.
7. Deselect Auto under Exposure and click microscope to manually adjust the LED intensity, Integration time, and Gain for optimal imaging settings. Auto exposure is not recommended for quantitative comparisons across wells. **Note:** Avoid high LED intensity to protect cells from phototoxicity and prevent photobleaching of fluorophores. Optimize the LED intensity and gain values for each channel using wells expected to have the highest and lowest fluorescence, such as cells that were treated with the highest or lowest concentrations of BSO. Too strong exposure will cause saturation of images. And too weak exposure may produce images with poor quality. Exposure settings that cover the whole range of heterogeneous imaging conditions are particularly important in a high throughput imaging context.
8. Designate a Fixed focal height according to the bottom elevation optimized in the plate definition step. Image the selected wells with the laser autofocus function which enables a whole 96-well plate to be imaged within 10 min when a single view field is imaged in each well. At a confluence of around 50% of Hela cells which were evenly distributed in wells, more than 1000 cells were imaged in each well.
9. The optimized imaging settings were applied to all samples to be imaged across plates in a same experiment. Save the experiment with captured images for data analysis.

##### Image processing

1. Open a stored experiment and click any images that have been captured.
2. Process images mathematically to subtract background and remove blurs caused by the optics system.
3. Perform Cellular Analysis using the RT-GSH adduct channel as the Detection Channel since it had a higher signal to background contrast compared to the RT channel.
4. Use Options to optimize analysis parameters as shown in Fig. 2. Set Threshold to 5000 to select the minimum pixel intensity value for the signal of interest. Set the minimum object size to 25 µm and the maximum object size to 100 µm to select cells and exclude debris or big cellular clumps. Object sizes should be selected depending on cell types. As a rule of thumb, it is recommended to first try different thresholds or object sizes and check if the expected cells are selected and then decide the final parameters.
5. ADD STEP to apply the analysis settings to all the images collected in the experiment.
6. Perform Ratio transformation to determine the GSH level which was calculated as the ratio of the Mean fluorescence intensity of RT-GSH to the Mean fluorescence intensity of RT of all the cells analyzed.

**Fig. 2.**
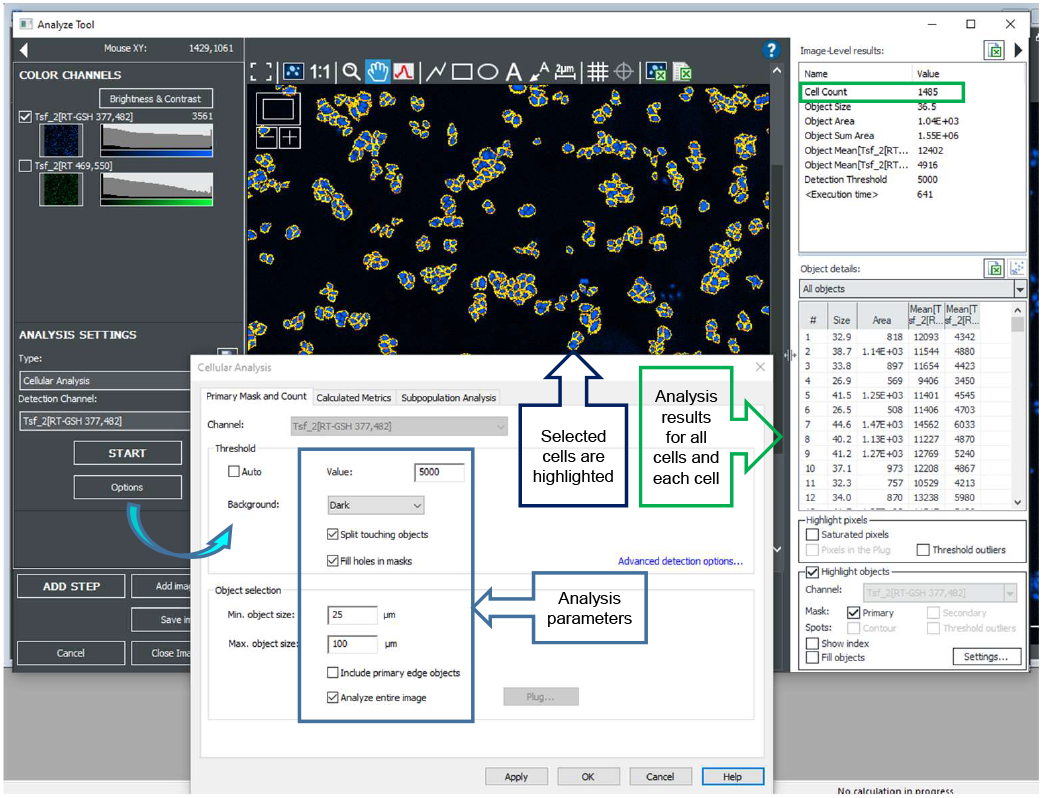
Cellular analysis setups. Select Options to set up the Threshold and the Min and max sizes for cells to be analyzed. The cells that meet the criteria were highlighted in the image window (middle of the screen). The analysis results for all the cells selected in the whole image were shown in the top right window and the analysis details for each cell were shown in the bottom right window.

#### 2.4.3 Data analysis

The data was analyzed and graphed using GraphPad Prism software. The ratios of the mean fluorescence intensity of RT-GSH to RT were averaged between the duplicates at each treatment condition and the standard deviations were calculated. The ratios of RT-GSH to RT were graphed against the logarithmic concentrations of BSO in Molar.

## 3. Representative Data and Discussion

γ-glutamylcysteine synthetase (γ-GCS) catalyzes the conjugation of cysteine with glutamate to form γ-glutamylcysteine which then conjugates with glycine to form glutathione[20]. BSO is a specific inhibitor of γ-GCS which has been observed lower glutathione concentrations *in vitro* and *in vivo*[21], [22]. As shown in Fig. 3a. GSH levels were decreased by BSO in a dose-dependent manner after 48 h of treatment, which is reflected by the decreasing mean fluorescence intensity ratio of RT-GSH to RT with increasing concentrations of BSO. GSH probe at the concentration of 2.0 µM accurately monitored the BSO treatment effect on GSH level in the living Hela cells. The mean fluorescence intensity reflected the average fluorescence intensity of RT-GSH and RT of more than 1000 cells that were imaged in each well of the two replicates treated with serial concentrations of BSO. The effect observed in such large cellular populations are expected to provide more comprehensive and less biased information than the conventional imaging approaches, such as using a confocal microscopy. This experiment was repeated more than 3 times, and it was well replicated with neglectable differences between replications. As shown in the representative images (Fig. 3b), GSH is distributed throughout the cells, including the cytoplasm and nuclei, and there were no significant morphological changes or death of cells were observed after imaging. In addition, the BSO treatment effect on GSH level was replicated in other cell lines, such as the human embryonic kidney (HEK) 293 cells and the microglial cell line BV-2 with the same concentrations of BSO and same concentration of RT probe using the same imaging system with some modifications of specific settings, such as LED intensity and Gain value (data not provided). Taken together, the present results demonstrated that the GSH probe RT sensitively and precisely monitored the GSH changes in the living Hela cells at a high throughput imaging platform and is expected to be applied to other cell lines.

**Fig. 3.**
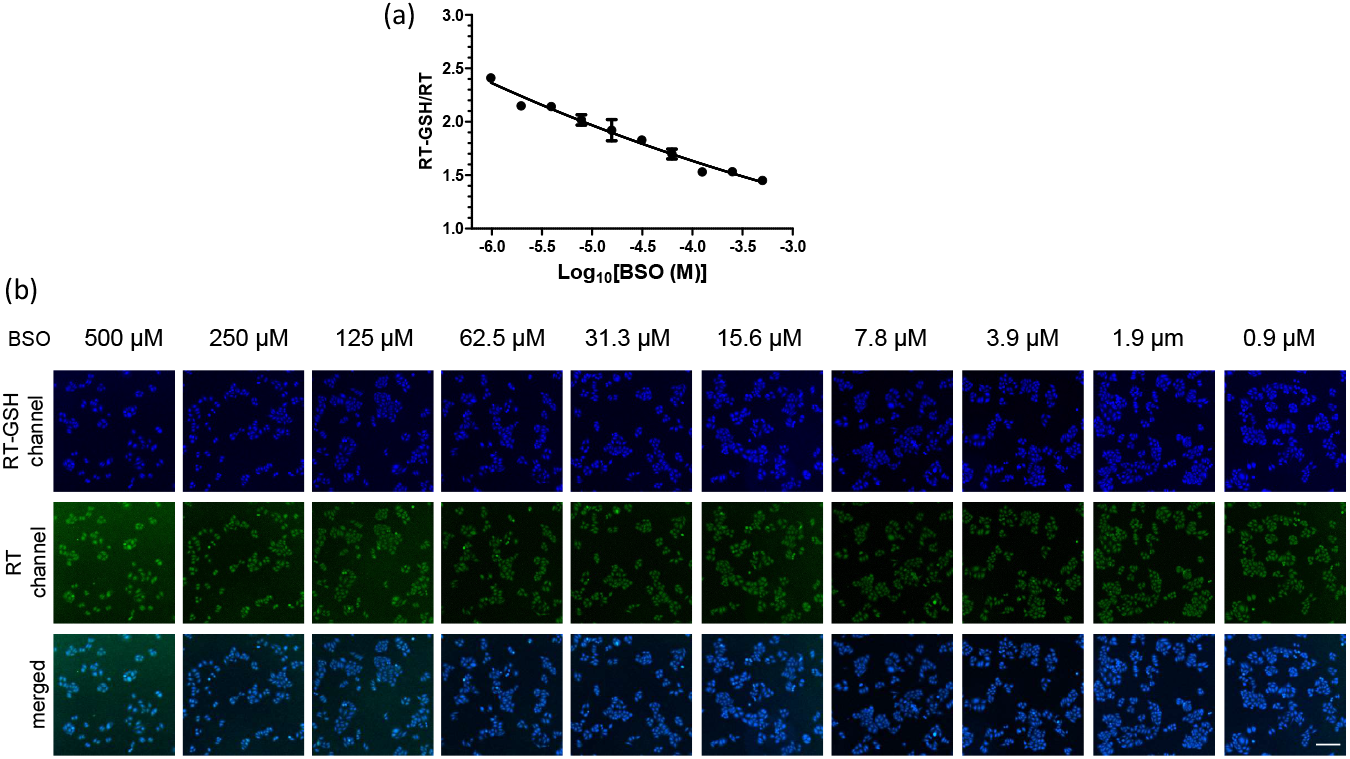
Quantification of GSH level after BSO treatment. (a) Quantitative analysis of GSH level in HeLa cells treated with serially diluted BSO (500 µM to 0.9 µM, 2 folds dilution) for 48 hours. Each data point represents the mean value and the standard deviation of 2 wells of cells in a 96-well plate. (b) Representative images of the dose-dependent changes of GSH levels in HeLa cells treated with serially diluted BSO for 48 hours. Scale bar, 200 µm.

## 4. Conclusions

The present work measured the GSH level with the GSH probe RT in Hela cells treated with varying concentrations of BSO using a high throughput live-cell imaging system. Images of more than 1000 cells from each well were captured and analyzed for fluorescence intensity of RT and RT-GSH, and the GSH level was reflected by the ratio of the mean fluorescence intensity of RT-GSH to RT. The results showed that GSH levels were decreased by BSO in a dose-dependent manner after 48 h of treatment, which is sensitively and precisely monitored by the RT probe. The present methods have the potential to be applied to any cell lines, although some specific imaging parameters may need to be optimized as different cells lines demonstrate varying concentrations of GSH levels and associated fluorescent properties.

## Acknowledgements

The research was supported in part by NIH grants R01-GM115622 and R01-GM115622-05S1 (to J.W.), Welch Foundation Q1912 (to M.C.W.), and Howard Hughes Medical Institute (to M.C.W.).

